# Very Early Responses to Colour Stimuli Detected in Prestriate Visual Cortex by Magnetoencephalography (MEG)

**DOI:** 10.1101/106047

**Authors:** Yoshihito Shigihara, Hideyuki Hoshi, Semir Zeki

## Abstract

Previous studies with the visual motion and form systems show that visual stimuli belonging to these categories trigger much earlier latency responses from the visual cortex than previously supposed and that the source of the earliest signals can be located in either the prestriate cortex or in both the striate (V1) *and* prestriate cortex. This is consistent with the known anatomical connections since, in addition to the classical retino-geniculo-striate cortex input, there are direct anatomical inputs from both the lateral geniculate nucleus and the pulvinar that reach the prestriate visual cortex without passing through V1. In pursuing our studies, we thought it especially interesting to study another cardinal visual attribute, namely colour, to learn whether colour stimuli also provoke very early responses, at less than 50 ms, from visual cortex. To address the question, we asked participants to view stimuli that changed in colour and used magneto-encephalography to detect very early responses (< 50 ms) in the occipital visual cortex. Our results show that coloured stimuli also provoke an early cortical response (M30), with an average peak time at 30.8 ms, thus bringing the colour system into line with the visual motion and form systems. We conclude that colour signals reach visual cortex, including prestriate visual cortex, earlier than previously supposed.

**Key points summary:**

- We measured visual evoked responses to colour stimuli using magnetoencephalography.
- An early response was identified, at around 30 ms after stimulation.
- The sources of the response were estimated to be in prestriate cortex.
- Colour signals thus appear to evoke very early cortical responses, just like form and motion signals.

**Abbreviations list:**

MEG: magnetoencephalography;
LGN: lateral geniculate nucleus;
EEG: electroencephalography;
ISI: inter-stimulus interval;
VEF: visual evoked magnetic field;
SNR: signal to noise ratio;
fT: femto Tesla;
ANOVA: analysis of variance;
MSP: multiple sparse priors;
MRI: magnetic resonance imaging;
RMS: root-mean-square;
SOI: sensor of interest.

## Introduction

It has long been known that the lateral geniculate nucleus (LGN), a thalamic station in the classical visual pathway from retina to cortex, projects not only to the primary visual cortex (striate cortex or area V1) but also to the visual areas of the prestriate cortex (Benevento & Yoshida, 1981; Fries, 1981; Yukie & Iwai, 1981; Gattass *et al*., 2014) and that visual signals are detected in the LGN in the 20–30 ms time window after stimulus onset (Maunsell & Gibson, 1992; Nowak *et al*., 1995; Givre *et al*., 1995). Electro-encephalographic (EEG) and magneto-encephalographic (MEG) studies have shown that signals related to two out of the three cardinal visual attributes, - forms (both simple and complex) and motion - can be detected in V1, V2 and V3 as well as in areas critical for the processing of face and house stimuli within the early time window of 25-45 ms (Shigihara & Zeki, 2013, 2014; Shigihara *et al*., 2016) while visual signals related to fast motion can be detected in V5 before V1, at around 30 ms (ffytche *et al*., 1995). This relatively new information about the organization of visual pathways to the cerebral cortex raises interesting questions about how the visual brain operates (see Zeki, 2016) but it leaves out of account another cardinal visual attribute, namely colour and whether signals related to it too can access the visual brain at early time frames through direct inputs as well as through ones relaying through V1. Previous studies have shown that the earliest responses to colour stimuli have a range of 72–120 ms (Paulus *et al*., 1984; Plendl *et al*., 1993; Schmolesky *et al*., 1998; Kuriki *et al*., 2005; Amano *et al*., 2006), considerably later than the responses elicited by form and motion stimuli (ffytche *et al*., 1995; Shigihara & Zeki, 2013, 2014; Shigihara *et al*., 2016). Given the early responses (at less than 50 ms) detected in LGN and in prestriate visual areas, we were interested to re-examine the time course of visual cortical responses to the viewing of coloured stimuli and to learn whether they, too, can be detected at early time windows, of 25–50 ms. The most obvious candidate for carrying colour-related visual signals to prestriate cortex, including V4, would be the classical visual pathway, from the LGN to V1 and then to V4, both directly and through V2 (Zeki, 1971; Zeki, 1978; Zeki & Shipp, 1989). But it is equally likely that a direct V1-bypassing route produced the responses we report here. Both the pulvinar and LGN are known to project directly to V4 (Cragg, 1969; Fries, 1981; Schmiedt *et al*., 2014; Yukie and Iwai, 1981) and this direct route is apparently sufficiently potent to enable a crude but conscious experience of colour (Morland *et al*., 1999), though the stimuli must be large (Brent *et al*., 1994).

## Material and Methods

### Participants and study design

Twenty four healthy adult volunteers (14 female, mean age 27.8 ± 5.4 years) took part in the study, and data from 18 participants (9 female, right-handed, mean age 28.3 ± 5.5 years) were analysed. Six participants were excluded from analysis, either because they slept in the scanner, had large body movement artefacts in the acquired data, performed poorly in the fixation task (see Results section for details), or because we encountered technical problems with their datasets. None of the participants had a history of neurological or psychiatric disorder; written informed consent was obtained from all and the study, which conforms to the Code of Ethics of the World Medical Association (Declaration of Helsinki; printed in the British Medical Journal 18 July 1964), was approved by the Ethics Committee of University College London.

### Stimuli and Task

Stimuli were generated using Cogent 2000 and Cogent Graphics toolboxes running in MATLAB (MathWorks, Na-tick, MA, USA). To stimulate the colour system preferentially and at the same time minimize the stimulation of the form system, we used colour stimuli with the following characteristics (Figure 1):

1. The stimuli consisted of three coloured circles (blue, red, and grey) (5.0° in diameter) but, in any given presentation, only two were displayed (e.g. blue and grey or red and grey or blue and red or grey and grey). The pairs of stimuli were displayed separately in either the lower left or lower right quadrants of the visual field (and subtended 2.5° - 7.5° in visual angle); this was done to avoid cancelation effects that can occur when both banks of the calcarine sulcus are stimulated (Portin *et al*., 1999).
2. The edges of the circles were blurred using sine curve filtering to minimize the stimulation of edge detectors in both V1 and V2; cells in V4 can also be sensitive to both edges and colours (Zeki, 1983; Bushnell & Pasupathy, 2012) though their orientation profiles are broader than equivalent cells in V1 and V2 (Zeki, 1983; Desimone & Schein, 1987).
3. Three different colours (blue, red and grey) were used because long-wave and short-wave inputs have significantly different anatomical distributions within the LGN (Martin *et al*., 1997; Chatterjee & Callaway, 2003; Mullen *et al*., 2008). The V1-bypassing input from the LGN to area V4 comes mainly from the intercalated layers (Lysakowski *et al*., 1988; Lyon & Rabideau, 2012), which is dominated by the short-wave cone input, while the parvocellular layers which project to V4 through V1 are dominated by long- and middle-wave cone inputs. It was therefore conceivable that blue and red stimuli would activate visual cortex differentially, both spatially and temporally. Hints that this may be so are found in the fact that red stimuli produce stronger responses in V1 and are more potent in eliciting seizures in photosensitive individuals than blue light (Givre *et al*., 1995; Takahashi & Tsukahara, 1998; Takahashi & Fujiwara, 2004).
4. We did not use a single intensity for all subjects; instead, we used the preferred method of varying the intensity to suit each subject individually, by using flicker photometry, as before (Bartels & Zeki, 2006: http://www.vislab.ucl.ac.uk/cogent.php). Hence, though the adjustments required differed from subject to subject, the intensities of the blue, red, and grey stimuli were adjusted to be perceptually equal to each other for each participant.

**Figure 1.**
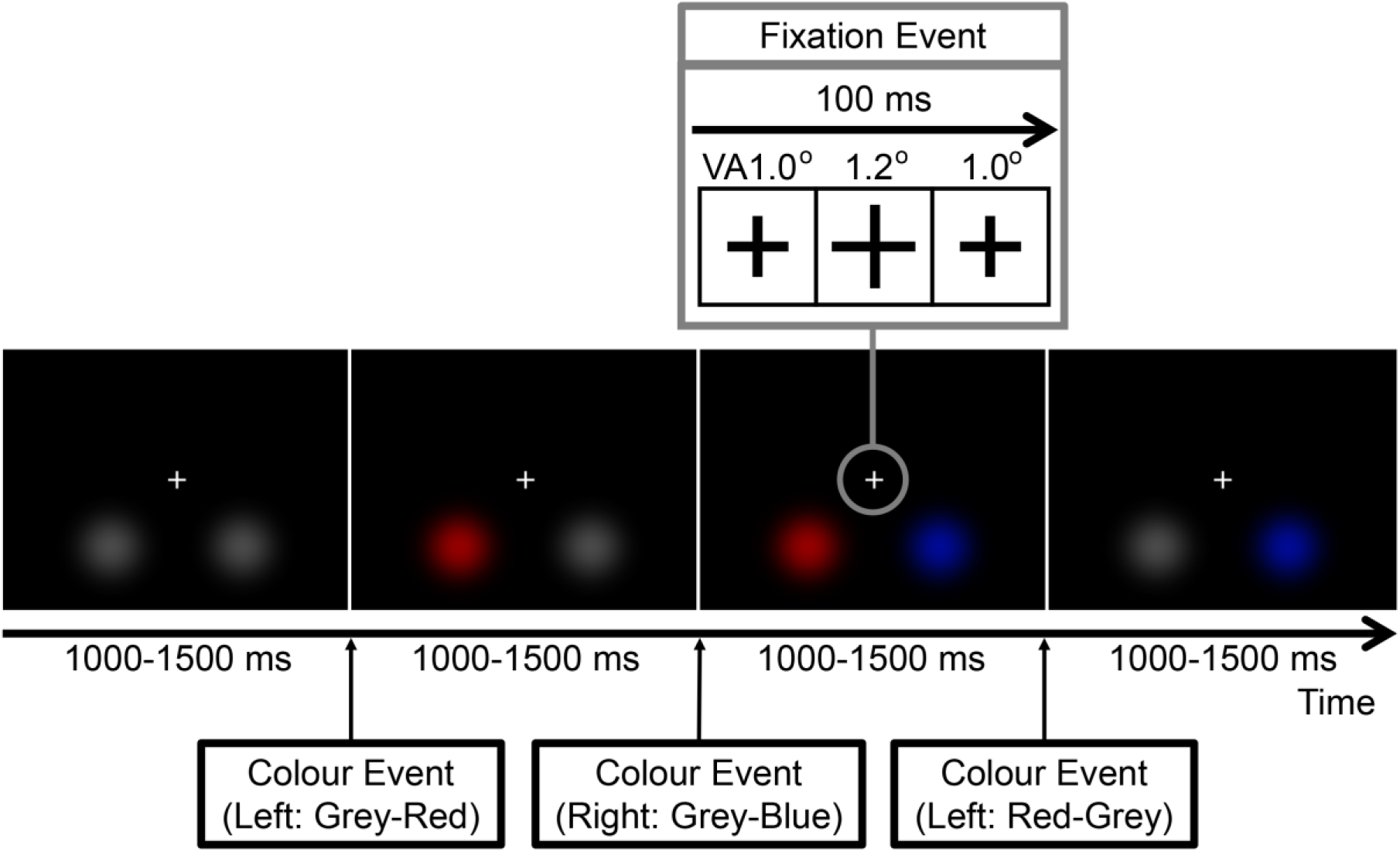
Schematic representation of the stimulation paradigm. There were 288 colour events per MEG scanning session, with random intervals between colour changes (1000-1500 ms). Within randomly selected 10 intervals, the fixation cross increased in dimensions (from 1.0 to 1.2°) for 100 ms (fixation event). Participants were instructed to keep fixating the cross and asked to press a button when the fixation event occurred. One MEG scanning session took 4 min 48 sec to 7 min 12 sec (varying according to the intervals); each participant performed 6 sessions with a short break between them.

Each participant had 5 to 6 scanning sessions in total. The duration of each session varied between 4 min 48 sec and 7 min 12 sec, depending on inter-stimulus intervals (ISI), which were randomly assigned at the beginning of each session. In a given session, participants viewed passively a change of the colour (colour event) in either the lower left or right quadrant with a randomly varied ISI of 1000–1500 ms. The colour event was a change in colour (grey to blue, red to blue, grey to red, blue to red, blue to grey and red to grey) which occurred in either left or right quadrant, amounting to a total of 288 colour events distributed in the two quadrants. The colour conditions (blue, red, and grey) were defined by the colour which appeared after the colour event (e.g. with blue, this was grey to blue or red to blue) and the order of the 12 changes was randomized within a session, during which participants were instructed to focus on a circular fixation point of (1.0° × 1.0°) located at the centre of the screen, and press a button when its size changed (fixation event). The fixation event occurred randomly 10 times per session, was 100 ms in duration and 0.2° in extent. The fixation events always occurred within the ISI of a colour event, so that the two events did not overlap temporally. A schematic representation of the paradigm used is given in Figure 1.

### Scanning Details

MEG scans were performed inside a magnetically shielded room. Participants were seated in the scanner and viewed a screen placed 60 cm in front of them. Stimuli (5.0° in diameter) were rear projected by a projector (DLA-SX21, Victor Company of Japan, Kanagawa, Japan) with a resolution of 800×600 pixels at 60 Hz. Trigger signals were recorded for the MEG system through an IEEE 1284 connection. We had determined through a photodiode placed on the screen that the delay between the trigger signal and the projection of stimuli was 33 ± 4 ms; during data processing, data were accordingly shifted by 33 ms, to correct for the delay. MEG data were recorded continuously using a 275-channel CTF Omega whole-head gradiometer (VSM MedTech, Coquitlam, Canada) and sampled at 1200 Hz with a 200 Hz low-pass filter. Participants were fitted with localizer coils at the nasion and 1 cm anterior to the left and right traguses to monitor head movement during the recording sessions.

### MEG data processing

Data were analysed offline using SPM-12 (Wellcome Trust Centre for Neuroimaging, London, UK). They were divided into 1000 ms epochs, each starting 500 ms before stimulus onset. Epochs in which the difference between the minimum and the maximum strength of the magnetic signal exceeded 10000fT were discarded by using peak2peak algorithm in SPM-12. To reject blink artefacts, the remaining epochs were decomposed by independent component analysis using Fieldtrip toolbox (Oostenveld *et al*., 2011) and FastICA, and the components which correlated with either of the three eye-tracking parameters (x-/y-movement and pupil diameter) with more than 0.2 in linear correlation coefficient were rejected (muted). The artefact-free epochs were averaged in each condition and baseline corrected. About 120 responses were recorded for each participant, condition and quadrant. The signal during the 100 ms period preceding stimulus onset was used as a baseline. Software filters produce artefacts (Acunzo *et al*., 2012; Ramkumar *et al*., 2013) and are not recommended (Vanrullen, 2011); like others before us (Noguchi *et al*., 2004; Inui & Kakigi, 2006; Acunzo *et al*., 2012; Shigihara & Zeki, 2014), we therefore analysed our results without filters to identify the early response. The pre-processed MEG data were analysed both at sensor- and source-levels, as described in detail in following subsections. Figure 2 shows the entire procedure schematically.

**Figure 2.**
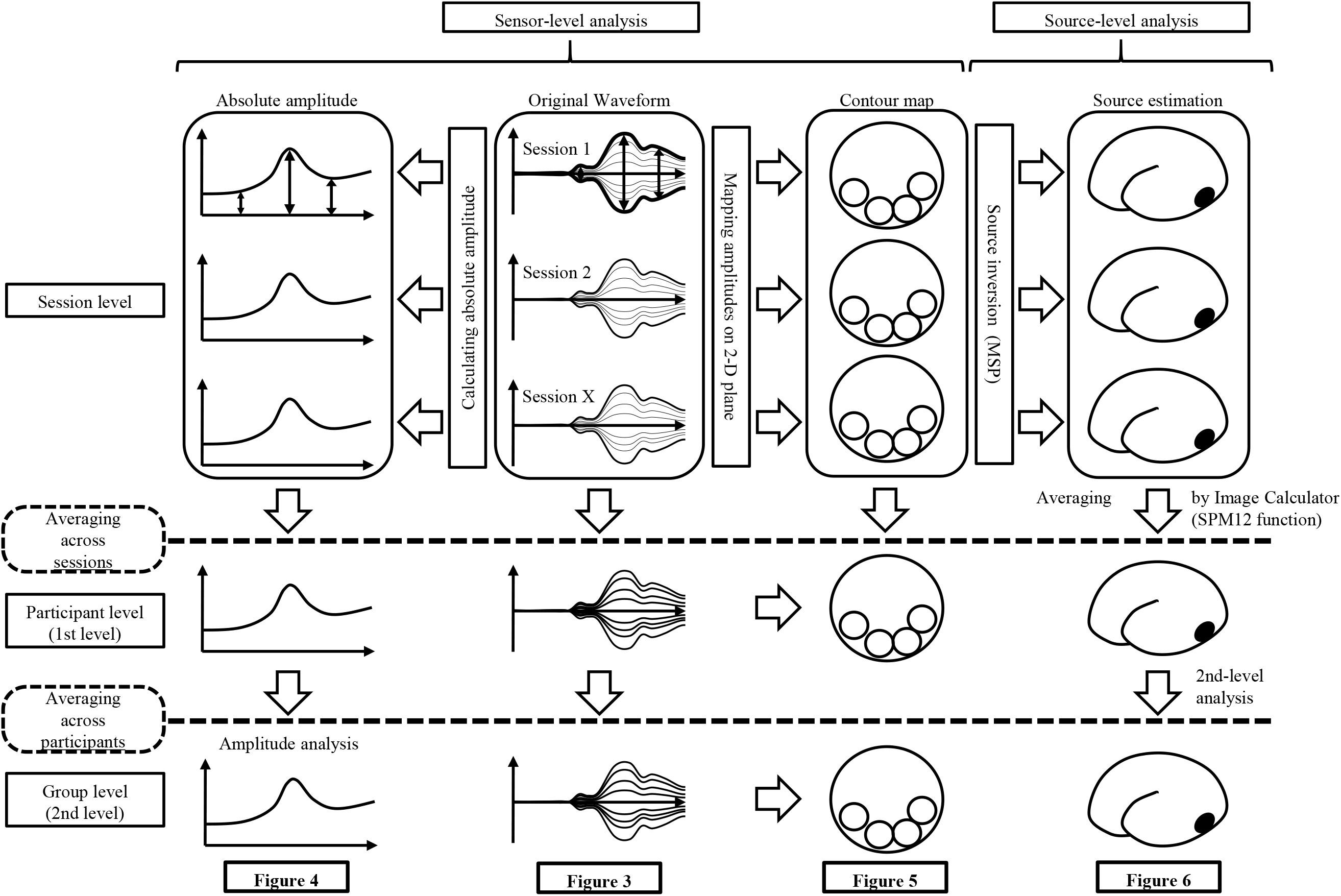
Schematic representation of the sequence of analysis. We started from the original waveform recorded in each session of each participant (middle upward in the figure). The sensor-level analysis consists of two steps: amplitude analysis (first panel (SOI Amplitude) and contour maps (third panel – contour map) analysis. In the amplitude analysis, we selected a sensor of interest (SOI) in each session for each participant, and averaged the amplitudes across sessions and participants (left hand side of the figure); we then compared the amplitude of the target time window against baseline. In the contour map analysis, we averaged out the original dataset between sessions and participants, and drew contour maps by mapping the sensor amplitude at the target time window on a 2D-plane (middle bottom in the figure). The source-level analysis was performed by inversing the source location from the sensor data using the Multiple Source Prior (MSP) algorithm in SPM8. The source locations were estimated for each session of each participant, and averaged out between sessions by Image Calculator function in SPM8 (single image was created for each participant – see right hand column). The averaged images were then taken to the 2nd-level analysis, and parametric maps were created (bottom right in the figure).

## Sensor-level analysis

### Data quality check

Sensor-level analysis is the central and essential part of this study; it provides the foundation for all other results, namely evidence that there is a very early response (< 50 ms) to colour events. Before starting the main part of the sensor-level analysis, we drew averaged visual evoked magnetic field (VEF) waveforms (butterfly maps) using a set of occipital sensors (MLO 11–53 and MRO 11–53; defined by SPM-12) for each quadrant presentation (lower left and right), to confirm that the recording and the pre-processing were properly undertaken (Figure 3).

**Figure 3.**
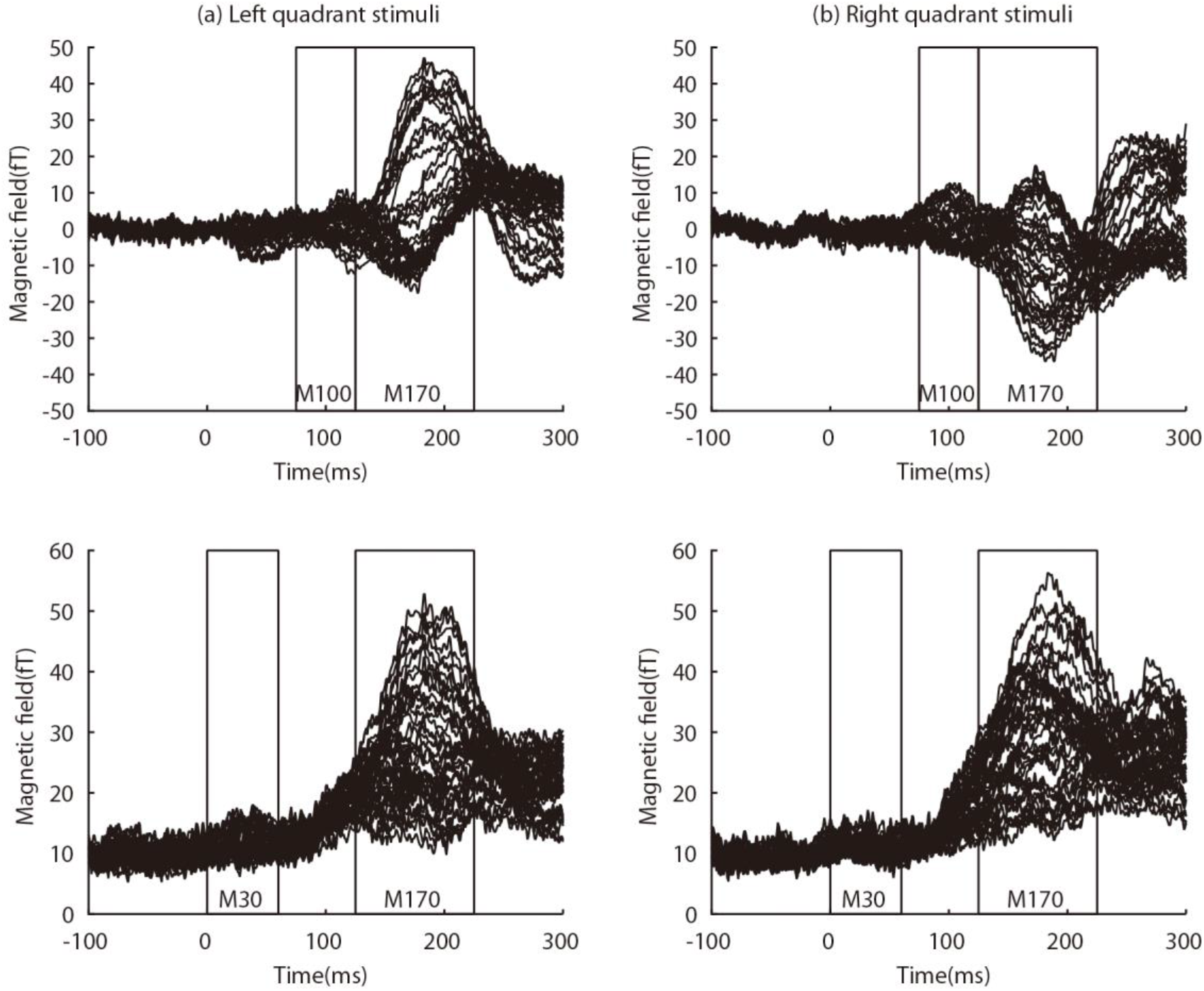
VEF waveforms measured by occipital sensors, averaged across 18 participants and both quadrants. The left figures (a) show the waveform resulting from left quadrant stimulation and the right figures (b) that resulting from right quadrant stimulation. The top figure in each column shows the raw waveform and the bottom figure does RMS waveform. Each of the 38 lines represents the averaged dataset in 38 occipital sensors. Two sequential responses, M100 and M170 are shown in the sections enclosed by rectangles in top figures. The polarities of the responses are opposite between left and right quadrant conditions. Although it is not as clear as the M100 and M170, M30 is also evident in the section enclosed by rectangles in RMS waveforms (bottom figures).

### Amplitude analysis

The amplitude analysis consisted of two steps: (1) visual inspection (identification) and (2) statistical analysis.

Although the VEF waveforms (butterfly maps: Figure 3) showed clear VEFs at around 100 (M100) and 170 ms (M170) after stimulus onset, responses at < 50 ms could not be identified visually because of their low signal to noise ratio (SNR). To identify them we used an absolute amplitude analysis, which is sensitive for detecting very early responses, as we had done in our previous study (Shigihara & Zeki, 2013). The absolute amplitude was calculated as the difference between maximum and minimum magnetic field intensities within a set of occipital sensors (MLO 11–53 and MRO 11–53; defined by SPM-12) at each sampling point. The left-most section of Figure 2 shows the calculated absolute amplitude waveforms schematically. For the visual inspection, they were averaged between all sessions and conditions, and plotted separately for each participant (Figure S1 in Supporting information). They were also averaged between all of the 18 participants, for each of the 6 conditions (3 colours × 2 quadrants: Figure 4a). To identify the average peak time by using the best SNR dataset, we also drew a waveform averaged across all conditions (Figure 4b), from which we defined the average peak time (30.8 ms).

**Figure 4.**
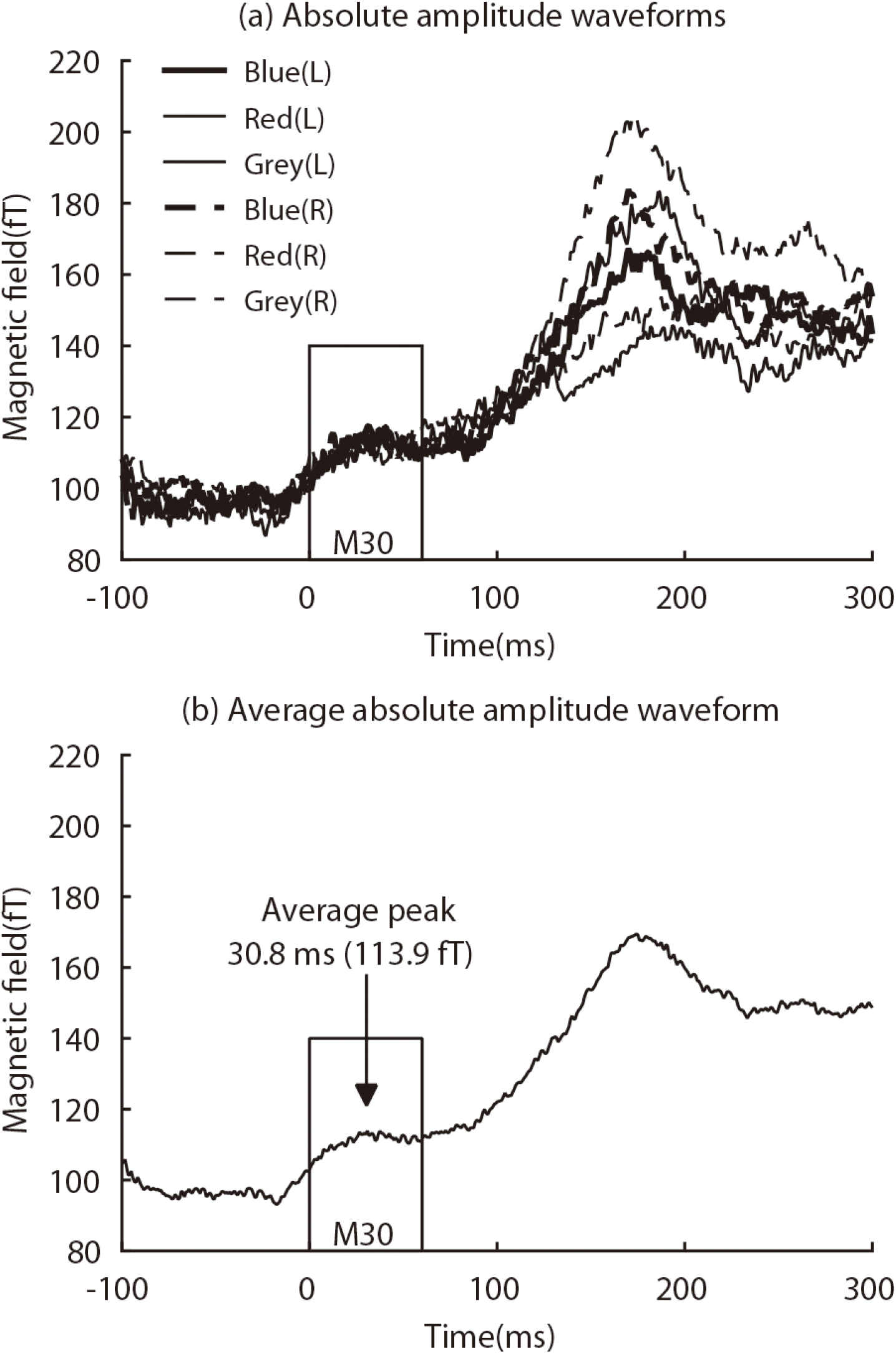
**(a)** Absolute amplitude waveforms averaged across 18 participants. Each of the 6 lines represents the averaged dataset in 6 conditions (blue, red and grey colour in lower left and right quadrant). Very early responses (M30) were obtained for all conditions (in the section enclosed by a rectangle). **(b)** Absolute amplitude waveforms averaged across 18 participants and all conditions, which also shows M30 responses with an average peak time at 30.8 ms after stimulus onset.

Next, statistical analyses were applied to confirm that early responses at less than 50 ms (M30) were not the result of baseline fluctuations but that they reflect significant VEFs. For each of the 6 conditions (3 colours × 2 quadrants), we compared the absolute amplitude at the peak time (30.8 ms) against the baseline amplitude (the average amplitude during 50 ms preceding the stimulus onset) by using paired t-test. Additionally, the amplitudes at the peak times were compared between the 6 conditions by a two-way ANOVA [3 colours (blue, red and grey) × 2 quadrants (left and right)] with repeated measures to which was applied a Greenhouse - Geisser correction (Greenhouse & Geisser, 1959) to adjust the degrees of freedom, if necessary. Prior to ANOVA, we confirmed the normality of the dataset, using Kolmogorov-Smirnov test (Massey Jr, 1951). The results of the statistical tests are shown in Table 1 and Table 2. Note that differences in peak latencies (peak times) were not compared (tested statistically) between conditions, while their amplitudes were. This is because of the difficulties and limitations of the statistical tests for peak times (see Discussion for details).

**Table 1.** The results of the statistical analysis (t-test) on the absolute amplitude responses (M30).

| Condition |  | Magnetic field amplitude (fT) |  |  |  | Paired *t*-test |  |
| --- | --- | --- | --- | --- | --- | --- | --- |
|  |  | Peak (30.8 ms) |  | Baseline (-50 to 0 ms) |  |  |  |
| Quadrant | Colour | Mean | *SD* | Mean | *SD* | *t* | *p* |
| Left | Blue | 117.4 | 34.4 | 97.4 | 24.8 | 6.55 | < 0.001 |
|  | Red | 115.9 | 37.5 | 95.0 | 21.8 | 3.89 | 0.001 |
|  | Grey | 110.7 | 26.6 | 95.0 | 23.0 | 5.35 | < 0.001 |
| Right | Blue | 113.8 | 31.6 | 97.9 | 22.3 | 4.51 | < 0.001 |
|  | Red | 114.1 | 25.1 | 98.7 | 24.5 | 7.03 | < 0.001 |
|  | Grey | 111.4 | 31.6 | 96.8 | 22.8 | 3.58 | 0.002 |

**Table 2.** The results of the statistical analysis (two-way ANOVA) on the absolute amplitude responses (M30).

|  |  | *F* | *df* | *p* |
| --- | --- | --- | --- | --- |
| Main effect | Colour | 1.19 | 2, 34 | > 0.1 |
|  | Side | 0.45 | 1, 17 | > 0.1 |
| Interaction |  | 0.14 | 1.4, 23.9 | > 0.1 |

### Contour-map analysis

Amplitude analysis showed that there was a significant M30 response; this was ascertained by studying the topographic patterns of the magnetic responses, using contour maps. Contour maps constitute a means of representing the sensor-level analysis to show the topographic pattern of the magnetic responses in two dimensions. They also reveal the orientation of the neural current (dipole orientation), which can help us to distinguish between two adjacent parts of the responsive cortex (Allison *et al*., 1991; Hämäläinen *et al*., 1993; Papadelis *et al*., 2011). Contour maps were drawn for each of the 6 conditions at the average peak time of M30 (30.8 ms) (Figure 5).

**Figure 5.**
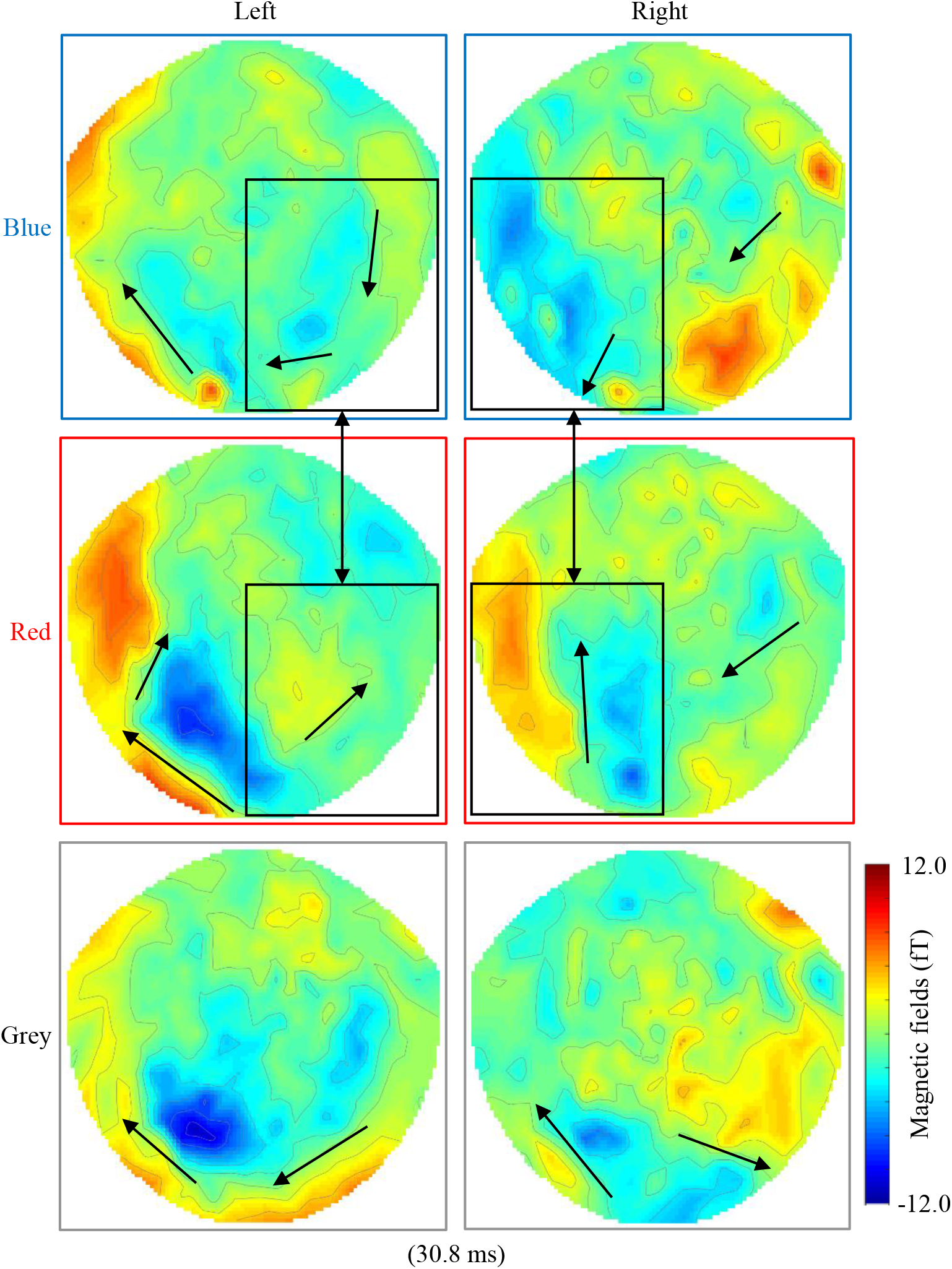
Contour maps drawn for each of the colour conditions in each quadrant at the peak time of M30 (30.8 ms). Note that all show at least two sets of combinations of inflow and outflow of magnetic fields in occipital sensors on both sides. Also note that, between blue and red colour conditions, estimated current orientations were different in the sections enclosed by rectangles (contralateral hemisphere to the stimulus presentation). Bold arrow, dipole orientation in occipital area.

### Source-level analysis

The two steps of the sensor-level analysis (amplitude analysis and contour-map analysis) described above showed that there was an M30 response to colour stimulation. Both (lateral) sides of the occipital areas were responsible for producing it (contour-map analysis). To estimate more precisely the locations of brain areas producing the response, we carried out a source-level analysis using Multiple Sparse Priors (MSP, Greedy search) (Mattout *et al*., 2005; Friston *et al*., 2008). Forward modelling was performed for the whole brain using a single sphere model (fine mode) with individual participant anatomical scans acquired by a 3T MRI scanner (Siemens Magnetron Allegra or Trio Tim 3T scanner, Siemens, Erlangen, Germany). Three out of 18 subjects could not participate in the structural MRI scans for a variety of reasons (tattoos, claustrophobia, etc.); we therefore used the standard brain template in SPM for them. The source inversion was performed for the 0–75 ms time window after stimulus onset, using MSP for each participant, session and condition separately. No source priors were used for source localization. The sources of the VEFs for the peak time of M30 (30.8 ms) were estimated by MSP without filters (session level analysis). These images were averaged for each participant using the Image Calculator in SPM-12 (first level analysis), which is a mathematical tool to manipulate sets of corresponding numbers in 3D space. The calculated images were taken to the second (group) level analysis using t-tests. Here we report the source locations of the peak responses at M30 corresponding to 30.8 ms, at a significance threshold of *p* (uncorrected) < 0.001. However, this was not essential because the sensor-level analysis had already confirmed that there were sources in the occipital cortex (Figure 4, 5 and Table 1). See also the right-most section of Figure 2, for a schematic representation of the series of source-level analyses.

## Results

### Behavioural results

To restrict the presentation of stimuli to the left and right lower visual fields, participants were instructed to fixate the central fixation cross throughout the MEG scanning sessions. To confirm that they were doing so, the central cross changed in size randomly (Fixation Event; 10 times in a session), which participants were asked to indicate by a button press. One participant performed poorly in the task (having a hit rate lower than 70%) and was therefore excluded from the following analyses. On average, in 87% (*SD* = 10.2) of total fixation events, they detected the events successfully and pressed the button within 2 sec after the event. As the change in the dimensions of the cross was very small (0.2°) and its duration short (100 ms) and as some participants reported difficulties in maintaining focus on the cross in a dark room, due to micro-saccades (reviewed in Martinez-Conde et al. 2004), the detection rate of 87% is a reasonable confirmation that fixation of the central cross was high for our purposes.

## Results from Sensor-level Analysis

### Data quality check

VEF waveforms for the left and right quadrant conditions are shown separately in Figure 3. Each of the 38 lines in the figures represents the averaged dataset in 38 occipital sensors. Two sequential late responses are evident with peaks around 100 ms (M100) and 170 ms (M170) after stimulus onset, (sections enclosed by rectangles in Figure 3). The former probably corresponds to the classical N75 or / and P100 response to visual stimuli, while the latter replicates the late responses to colour stimuli reported by others (Plendl *et al*., 1993; Kuriki *et al*., 2005). The polarities of the M100 and M170 responses are opposite between the two presentation quadrants (lower left and right). This shows that the MEG recording and analyses were done properly. These late responses (M100 and M170) are not, however, the main interest of the present study, which is concerned with the very early responses after colour events; we therefore did not study them further.

### Amplitude analysis

To obtain an effective detection power for the very early component of the VEF, the absolute amplitude waveforms were calculated for each of the 6 conditions (Figure 4a). There are clear responses with peaks at around 30 ms after stimulus onset (M30) for all conditions (section enclosed by a rectangle in Figure 4a). Figure 4b shows the absolute amplitude waveform averaged across all conditions, which has the best SNR available in the present study. It also shows clear M30 response (section enclosed by a rectangle) with a peak time at 30.8 ms after stimulus onset. For detailed inspection, the absolute amplitude waveforms were separately drawn for each participant (Figure S1 in Supporting information). Individual waveforms showed clear peaks, which correspond temporally to M30 (at around 30 ms after stimulus onset): this is indicated by black arrows in the Figure S1 in Supporting information, for 13 out of the18 participants.

For each condition, the magnetic amplitude at the peak time (30.8 ms) was compared against the amplitude during the baseline period (-50 to 0 ms before stimulus onset) using paired *t*-test. The results were significant for all conditions, indicating that all of the 6 conditions showed the significant M30 responses. We also compared the peak amplitudes between the 6 conditions using two-way ANOVA with repeated measures. None of the main-effects of the quadrant or colour conditions, or the interactions, were significant, indicating that the peak amplitudes of M30 responses were not significantly different between the conditions or hemispheres. The results of those statistical tests are shown in Table 1 and Table 2.

### Contour-map analysis

Contour maps for each of the 6 conditions (3 colours × 2 quadrants) at the peak time of M30 (30.8 ms) are given in Figure 5. All show at least two sets of combinations of inflow and outflow of magnetic fields in occipital sensors on both sides, indicating that there was at least one brain area on each side producing the response. The topographic pattern was not the typical one for a magnetic source produced solely by V1 (Seki *et al*., 1996; Hatanaka *et al*., 1997; Shigeto *et al*., 1998). Furthermore, the patterns of magnetic fields indicate that the orientations of the currents produced by the neural responses, when the stimuli were presented in the contralateral quadrant, were different for the blue and red conditions (section enclosed by rectangles in Figure 5). This suggests that different colours produce responses in different, though adjacent, zones within a brain area located in prestriate cortex (see Discussion for details).

### Results from source-level analysis

We estimated the source of the VEF components at the peak time of M30 (30.8 ms after stimulus onset), in which significant magnetic responses were found in the amplitude analysis and the responses in occipital areas were confirmed through the contour-map analysis. To learn which brain areas were the most likely to have produced the response, we estimated the source of the signals using MEG at 30.8 ms after stimulus onset. Estimated source locations for the 6 conditions are shown in Figure 6; they were estimated to have been in visual cortex, and mainly in prestriate cortex, including area V4 (colour area: indicated by yellow circles in Figure 6). This suggests that our analyses detected the M30 response correctly. The results of source estimation did not show explicit differences in the locations of estimated magnetic source between conditions, although the estimated current orientation in the contour maps (section enclosed by rectangles in Figure 5) is different between conditions (see Discussion).

**Figure 6.**
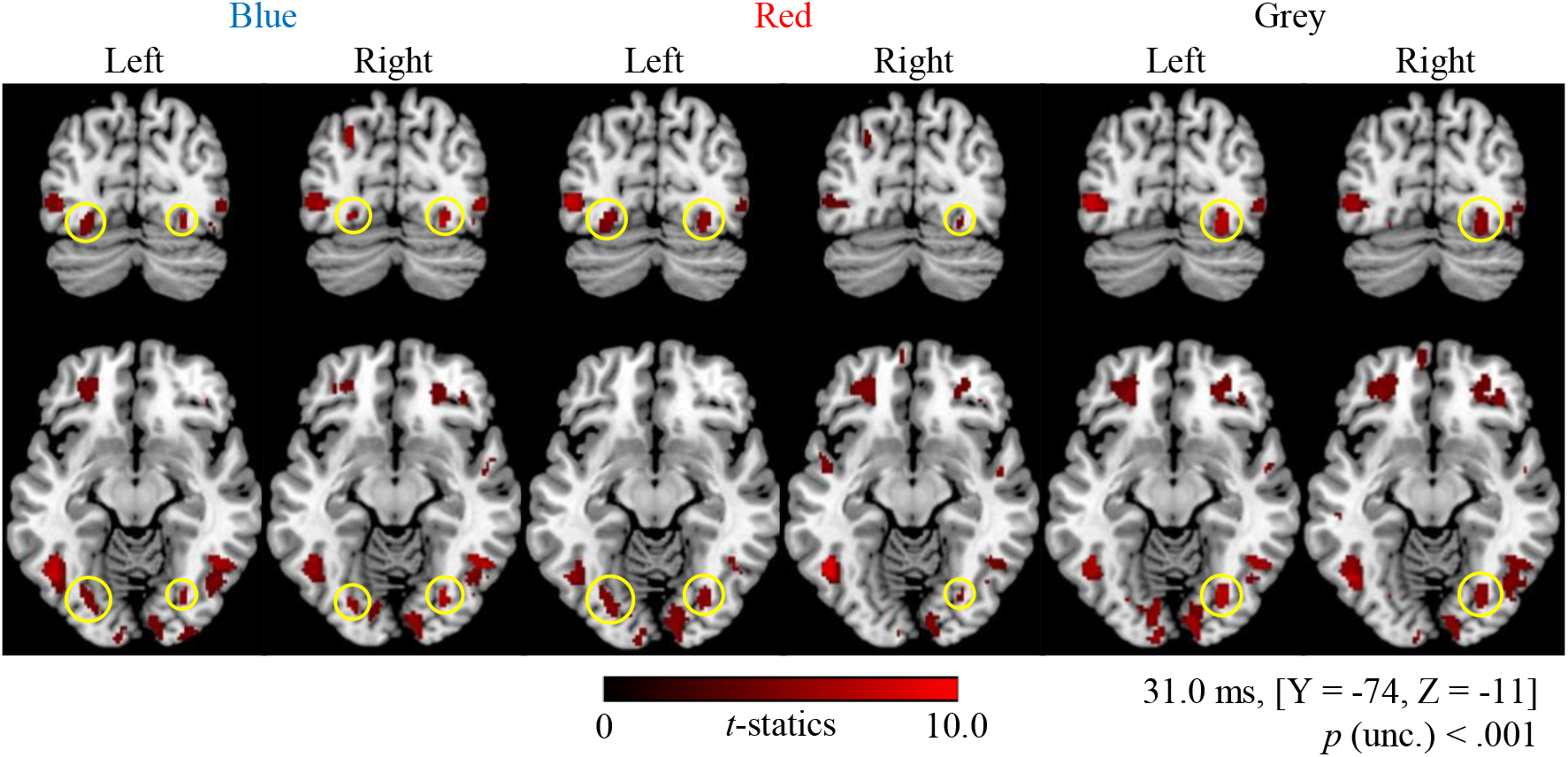
Statistical parametric maps estimated for each colour condition in each quadrant at 30.8 ms (M30), which is the earliest time window at which we detected responses. The display threshold is at peak level *p* (uncorrected) < .001, and the clusters in the V4 complex are identified by yellow circles.

Summarising these results, our sensor-level analysis showed that the very early response with our colour stimuli had an average peak at 30.8 ms (M30) ms after stimulus onset, that this response was produced mainly by prestriate cortex (including area V4) and that different coloured stimuli yield responses from adjacent parts of this active zone (as detected by dipole orientation in contour maps).

## Discussion

Our principle finding in this study is that the viewing of coloured stimuli produces a very early response (M30) in the visual brain, with a peak at 30.8 ms after stimulus onset. We demonstrated this by (absolute) amplitude analysis, using approach established in our previous studies (Shigihara & Zeki, 2013, 2014; Shigihara *et al*., 2016). All 6 conditions constituting the experiment were analysed independently and showed similar VEF waves with a peak at around 30.8 ms as average (Figure 4a), close to the time window of the very early response for fast motion (about 30–35 ms) (ffytche *et al*., 1995), for form (27–42 ms) (Shigihara & Zeki, 2013) and for checkerboard patterns (25–30 ms) (Shigihara *et al*., 2016).

By using the same MEG technique to show that colour stimuli elicit a response at about 30 ms from visual cortex (M30), this demonstration brings colour into line with form and motion, both of which provoke significantly earlier responses from visual cortex, including prestriate visual cortex, than previously supposed.

### Technical advantages of the present study

The average peak of the M30 response to colour stimuli was found to occur at 30.8 ms after stimulus onset and was earlier than the ones reported in previous studies (72–120 ms) (Paulus *et al*., 1986; Schmolesky *et al*., 1998; Kuriki *et al*., 2005; Amano, 2006). There are two reasons which may account for our success in detecting these very early responses, which are not evident in past studies: (1) We used absolute amplitude analysis in which the M30 was clearly identified in VEF waveforms (Figure 3). Since the M30 was so clear (see Figure 3), we did not need to use absolute amplitude analysis, although we did so to confirm our findings. Earlier studies may have ignored this early component and only focused on salient later responses (e.g. M100, M170) in the VEF waveforms. Through our earlier studies (Shigihara & Zeki, 2013, 2014; Shigihara *et al*., 2016), we have established protocols to detect small, early, responses; this consists of an absolute amplitude and sensor of interest (SOI) analysis, together with affiliated statistical analyses, followed by a contour map analysis. As described in the Methods section, the amplitude analysis is sensitive for detecting weak responses, evoked in local areas of the brain and recorded with a small number of sensors. The dynamic difference in the scales of the magnetic amplitudes between VEFs (Figure 3: within the range of - 50 to 50 fT) and absolute amplitude waveforms (Figure 4: within the range of 120 to 270 fT) reflects the difference of the sensitivity between the two methods. In the standard VEF waveform (Figure 3), the averaging of the waveforms across the sessions, participants, and conditions reduces the absolute amplitudes of the waveforms. The same sensor may record the responses with opposite polarity between sessions (due to the difference of the head position) and between conditions (due to the difference of the cortical source). Therefore, averaging across the datasets with such opposite polarities (positive and negative values) would potentially cancel out the amplitudes in the resulting waveforms. The absolute amplitude of the waveforms (Figure 4) do not suffer from the cancellation effects, since there are no negative values. Our tailored analysis method to detect the small response thus enabled us to identify the M30 response. (2) We used isoluminant colour stimuli with blurred edges for stimulating the lower quadrants: such stimuli activate the colour system preferentially and reduce activation of the form system, hence lessening any contamination between the two systems and maximizing the sensitivity of the MEG recording (Leventhal *et al*., 1998; Song & Baker, 2007).

### Reliability of detecting very early responses to colour stimuli

Although the absolute amplitude waveforms showed clear M30 responses with the peak around 30.8 ms post stimulus (Figure 4), we wanted additional evidence to validate the finding. There are three lines of evidence: (1) significant responses shown through amplitude analysis; (2) a difference in dipole orientation between blue and red stimuli (section 3.2.3); and (3) a reasonable and expected source estimation in prestriate cortex, including the colour area (V4). These results were independent of each other and all strongly support our main conclusion - that signals activate the visual brain much earlier than previously supposed. The M30 responses to the colour stimuli at around 30.8 ms post stimulation that we report are, therefore, unlikely to be the consequence of noise or artefacts obtained through inappropriate analysis. It is perhaps worth noting that other methods for analysing our results, though less satisfactory, also give the same result - of a very early (M30) response (see Supporting information).

In summary, the present work identified very early magnetic responses (M30) to colour stimulation, with the average peak at 30.8 ms after stimulus onset, and this response was derived from occipital areas, mainly from prestriate cortex. This conclusion is supported by relatively reliable evidence. When considered with evidence from the “estimation” approach, it points to V4 as one of the brain areas producing magnetic responses to colour stimuli at the early time window of < 50 ms. Nor is our study the only one to show such an early response. Although they do not mention or discuss it at all, Figure 3B and C (and especially 3B) of the paper by Bartsch *et al*. (2017) on cortical responses following attention to colour using MEG show clear early responses; from their figure, we calculate that these early responses occurred at about 45 ms. It is a pity that this early response, though evident in their figure, is not considered further. It merits study for, if confirmed as an early response, it would lead to a significant re-interpretation of the direction of traffic flow in the cortex which they outline in their paper. We are therefore led to the following conservative conclusion: that the M30 response is due predominantly to activity in prestriate cortex, including area V4. We note that we are not able to exclude early activation in V1 from our data. It would not be surprising if V1 were also co-active at this early time window, since the results we obtained previously using face and house stimuli show an early activation of V1 along with the prestriate cortex [Shigihara and Zeki, 2014]. In addition, the activation may have as well included (and probably did) areas allied to V4, such as V4α which, together with V4, constitute the V4 complex (Bartels & Zeki, 2000).

## Impact on physiology

### Pathways for transmitting rapid signals

We have no direct evidence to decide which pathway(s) were used to produce the M30 responses in our study, and hence do not wish to be dogmatic in our interpretation. Our reason for emphasizing the possibility that a V1 by-passing pathway may have delivered early signals to V4 lies partly in the fact that the sensor-level and contour-map analyses both point to prestriate cortex (including V4) as the source of the M30 responses and partly because this interpretation would be consistent with the results obtained for the brain’s motion and form systems (ffytche *et al*., 1995; Gaglianese *et al*., 2012; Shigihara & Zeki, 2013), and in particular the former, in which it has been shown that signals from fast-moving stimuli (>20° sec^-1^) result in responses from V5 before V1 (Beckers & Zeki, 1995; ffytche *et al*., 1995).

### Differences between colours

There were 6 conditions in the present study: 3 colours (blue, red and grey) × 2 quadrants (lower left and right), and M30 responses were detected for all (Figure 4 and Table 1). The consistency among all conditions strongly supports the existence of the M30 response to colour.

Past evidence shows some anatomical and clinical differences related to blue and red (dominated by short- and long-wave inputs, respectively). Short-wave (S) dominated cells occur mainly in the intercalated layers of the LGN which project directly to V4 without passing through V1 (Lysakowski *et al*., 1988; Lyon & Rabideau, 2012). On the other hand, red is more potent than blue in activating cells in V1 (Givre *et al*., 1995) and in precipitating seizures in photosensitive individuals (Takahashi & Tsukahara, 1998; Takahashi & Fujiwara, 2004). While we could not detect any difference in the amplitude of activation between blue and red light, we detected spatial differences in the sources of the early magnetic signals for these two colours. In the contour map analysis, the magnetic fields showed different patterns in the occipital areas contralateral to the presented stimuli (sections enclose by squares in Figure 5), and estimated current orientations from the magnetic patterns clearly have different, very nearly orthogonal directions (black arrows in Figure 5) between blue and red conditions. This would seem to indicate that the actual current locations have different normal directions in the cortex between the two conditions (e.g. located in different banks of the same sulcus). Previous studies using somatosensory responses found very tiny (< 1cm) locational differences in the somatosensory cortex, distinguished from each other through dipole orientation (Allison *et al*., 1991; Hämäläinen *et al*., 1993; Papadelis *et al*., 2011). Since, in our study, the principal source location for the early time frame was estimated in prestriate cortex and more particularly in V4, the difference in occipital current orientations within that early time frame implies that, likewise, the magnetic sources for the different colours (blue and red) had different locations within the responsive area, which we presume to be V4. It is known that neurons responsive to different colours are clustered within V4, in both monkey and humans (Zeki, 1977; Conway & Tsao, 2009; Brouwer & Heeger, 2013). Our results mirror these findings, and indicate that clusters within V4 responding to different colours within the early time frame are segregated from each other. Such small spatial differences in the cortex cannot be detected by the source inversion algorithm used in MEG analyses, because such analyses are based on the absolute amplitudes of the magnetic responses and do not take into account the orientation of the current. Even fMRI studies would be hard put to discriminate adjacent clusters responding to different categories of an attribute within a visual area. Our study therefore constitutes a good example of the importance of information derived from current orientation in neuroimaging studies using MEG.

Of course, the conclusions drawn from our present study are limited to the stimuli we used; they may or may not have been optimal and other stimuli may result in faster or slower delivery of signals to, and activity in, the visual brain.

### Limitations

There are five limitations to this study, which we discuss below:

Firstly, we did not compare the peak latency of the M30 responses between conditions. To perform a statistical comparison on latencies, the latency for each individual and for each condition would be required. This was not practical because of the low SNR: amplitudes of M30 were small and not using filters prevents us from improving the SNR. We therefore restricted ourselves to identifying peak times at a group level. This, however, does not affect our conclusion that there is an average peak at around 30 ms.

Secondly, there is a ± 4 ms jitter in timing of stimulus onset on the screen. Although this renders the SNR lower, the very early response (M30) was clearly identified in spite of it. The fact that the early response could be identified in spite of the jitter and the consequent lowering of the SNR, gives us added confidence in our results.

Thirdly, there is the possibility that the results were contaminated by signals from the retina and the LGN. For reasons given in Shigihara et al. (2016), we think that the retina and LGN cannot be the sole sources of the M30 response. Figure 5 shows some magnetic responses in the front part of the head: this might have had its origin from either inside or outside the brain. We have already shown that retinal neural activity affects MEG recordings and this could have contaminated the data (Shigihara *et al*., 2016). We took account of this possibility and did not use source priors to estimate sources either inside the visual brain or outside it at the single subject level, thus making our second level analysis free from contamination by these factors.

Fourthly, there are two established approaches for detecting very early responses: absolute amplitude analysis (Shigihara & Zeki, 2013) and SOI analysis (Liu *et al*., 2002; Noguchi *et al*., 2004; Shigihara & Zeki, 2014; Shigihara *et al*., 2016). We have used absolute analysis here, but we have also confirmed that similar results are obtained by using the SOI approach (See Supporting information).

Fifthly, there are two possible confounding factors to the early responses, which are micro-saccades and bottom-up attention. However, we expect that the micro-saccadic artefacts should be cancelled out or minimised, since we have averaged out the MEG signals over 100 trials participants. Additionally, bottom-up attention influences later components (e.g., M100) but there is no any evidence that it contaminates the very early cortical responses that we have addressed in the present study (M30). Therefore, we do not believe that the confounding factors influenced on the early responses significantly.

## Conclusion

In this work, we have demonstrated a very early response (M30) following colour stimulation derived mainly from prestriate cortex, including area V4. This brings the colour system into line with two other cardinal visual systems, namely those critical for form and motion perception, in showing that all three systems (colour, form, and visual motion) activate the visual brain within an early time frame. This is presumably done both directly, through the V1-bypassing systems, as well as by pathways using V1 to reach their destinations. This raises a host of interesting issues, which we have partially addressed (Zeki 2015, 2016) and which we will address more extensively in the future.

## Supporting information

The supporting information is available in the online version of this article.

